# Dynamic emotional updating as a computational marker of well-being

**DOI:** 10.64898/2026.02.05.703205

**Authors:** Noa Nagar, Kirill Vasilchenko, David C Jangraw, Robb B Rutledge, Hanna Keren

## Abstract

Mood shapes behavior, physiology, and well-being. While mood continuously fluctuates in response to external events, it can also remain surprisingly stable, raising a fundamental question: what shapes emotional well-being, mood stability or variability? Quantifying mood along these two dimensions is challenging. We combined a closed-loop mood modulating task with computational modeling to derive quantitative markers of emotional stability and variability and map them onto the continuum from well-being to depressive symptoms. Participants (n=209) experienced adaptive reward-based mood modulation and modeling revealed that higher well-being associates with mood variability and stronger weighting of recent events, whereas depressive symptoms were associated with greater mood stability and stronger anchoring to earlier events. Results replicated in an independent test–retest dataset, which also showed that these mood updating parameters are reliable across days. Our findings identify adaptive responsiveness to recent experiences as a quantitative marker of well-being and unravel the temporal mood dynamics in health and depression.

## Introduction

Well-being reflects a person’s overall mental health and is characterized by complex dynamics, with changes unfolding across multiple time scales. These dynamics emerge from various interacting processes, cognitive, emotional, and physiological, that integrate past experiences and current events to sustain or modify mood. This balance between mood stability and variability is a core dimension of mental health and understanding how it is disrupted in depression remains a fundamental question.

While daily life events often trigger mood shifts, individuals may also exhibit mood stability, whereby they return to their baseline well-being over time even after major life events (Brickman et al., 1978; Diener et al., 1999; Lucas et al., 2003; Wu, 2001, Rutledge et al., 2014, Houben et al., 2015). The ability to sustain positive mood states in the face of stress or difficult life events, has been linked to emotional resilience and optimal mental health (Diener et al, 2018; Tugade & Fredrickson, 2004). At the same time, emotional variability – the ability to dynamically adjust one’s mood in response to environmental demands – has been also proposed to be essential for well-being and resilience (Rottenberg et al., 2005; Waugh et al., 2011; Houben et al., 2015). Importantly, both extremes – excessive mood stability and excessive mood variability – can be maladaptive (Rottenberg et al., 2005; Waugh et al., 2011; Koval et al., 2013; Houben et al., 2015; Eldar et al., 2016). Formalizing the balance between mood stability and variability, and how it differs across individuals, can provide insights into the mechanisms of well-being and depression.

Imagine two people who face a minor setback at the beginning of their day: missing the bus to work. One reacts with frustration, but soon redirects focus, finds an alternative route, and moves on – returning to a good mood when the next positive event occurs. The other remains stuck in frustration and remains in a low mood for hours, regardless of later positive events, which colors their interactions, decisions, and productivity for the rest of the day. There are fundamental differences in how people update their mood in response to the environment and whether they can flexibly adjust to current circumstances, or whether early experiences persist and dominate.

To experimentally study well-being and mood, many works are focusing on self-reported mood valence as an index of well-being (Barrett, 2006; Rutledge et al., 2014; Keren et al., 2021; Liuzzi et al., 2022; Jangraw et al., 2023). Such mood ratings have been indeed shown to strongly correlate to depression symptoms across both clinical and subclinical populations (Tugade & Fredrickson, 2004; Fujita & Diener, 2005; Keyes, 2005; Rottenberg et al., 2005; Houben et al., 2015; Rutledge et al., 2017; Jangraw et al., 2023). However, studying mood dynamics and mood transitions is limited in several ways. Traditional mood-induction paradigms using stimuli such as emotional pictures, film clips, music, or written prompts (e.g., Gilet & Jallais, 2010; Zhang et al., 2014; Uhrig et al., 2016; Siedlecka & Denson, 2019). While such stimuli do not vary in intensity, monetary reward-based stimuli offer quantitative parametric scalability. Moreover, typical behavioral paradigms are open-loop – that is, the stimuli are pre-defined and not adaptive to participants’ ongoing changes in emotional responsiveness. This limits the ability to induce ongoing and robust mood transitions.

Thus, we developed a closed-loop mood modulating task that uses monetary wins and losses to modify participants’ mood in a personalized and adaptive manner.

This task builds on a probabilistic decision-making paradigm (Rutledge et al., 2014), where on each trial participants choose between a certain monetary amount or to gamble between two values, and experience various reward-prediction-errors (RPEs). This task engages value-based computations, anticipation processes, and rapid momentary mood responses (Mellers et al., 1997; Rutledge et al., 2014; Eldar et al., 2016; Keren et al., 2021; Liuzzi et al., 2022). The task’s strength lies in its ability to elicit mood fluctuations. By embedding a closed-loop circuit within this framework, our mood modulation utilizes this cognitive-emotional coupling while enhancing experimental control and efficacy and minimizing habituation. Importantly, it can allow us to compare mood variability versus stability in different settings and both within and across sessions.

We implemented this paradigm to shift and stabilize mood to either high or low states, as well as to three alternating mood states (high–low–high) on two consecutive days, enabling us to test reproducibility and reliability across time.

To parametrically model mood dynamics and quantify the parameters of updating versus rigidity of mood, we used the Primacy Mood Model (Keren et al., 2021), an extension of the Happiness Model by Rutledge et al. (2014), featuring two key temporal parameters: a recency coefficient (*β*_*R*_), reflecting responsiveness to most recent outcomes, and a primacy coefficient (*β*_*P*_), capturing the lasting impact of earlier outcomes. These parameters allowed us to quantify individual mood updating patterns ranging from recency-weighted adaptivity to primacy-anchored stability, and uncovering computational trait-like markers of emotional dynamics and their relation to depression.

## Results

### Differences in mood variability between healthy and depressive participants

To study individual differences in mood variability versus stability, we used a probabilistic monetary rewards task that provides different Reward Prediction Errors (RPEs) values, previously shown to influence momentary mood ratings (Rutledge et al., 2014). RPEs were dynamically adjusted based on participants’ ongoing mood reports, to maximally influence mood in an individualized manner (Figure 1).

**Figure 1.**
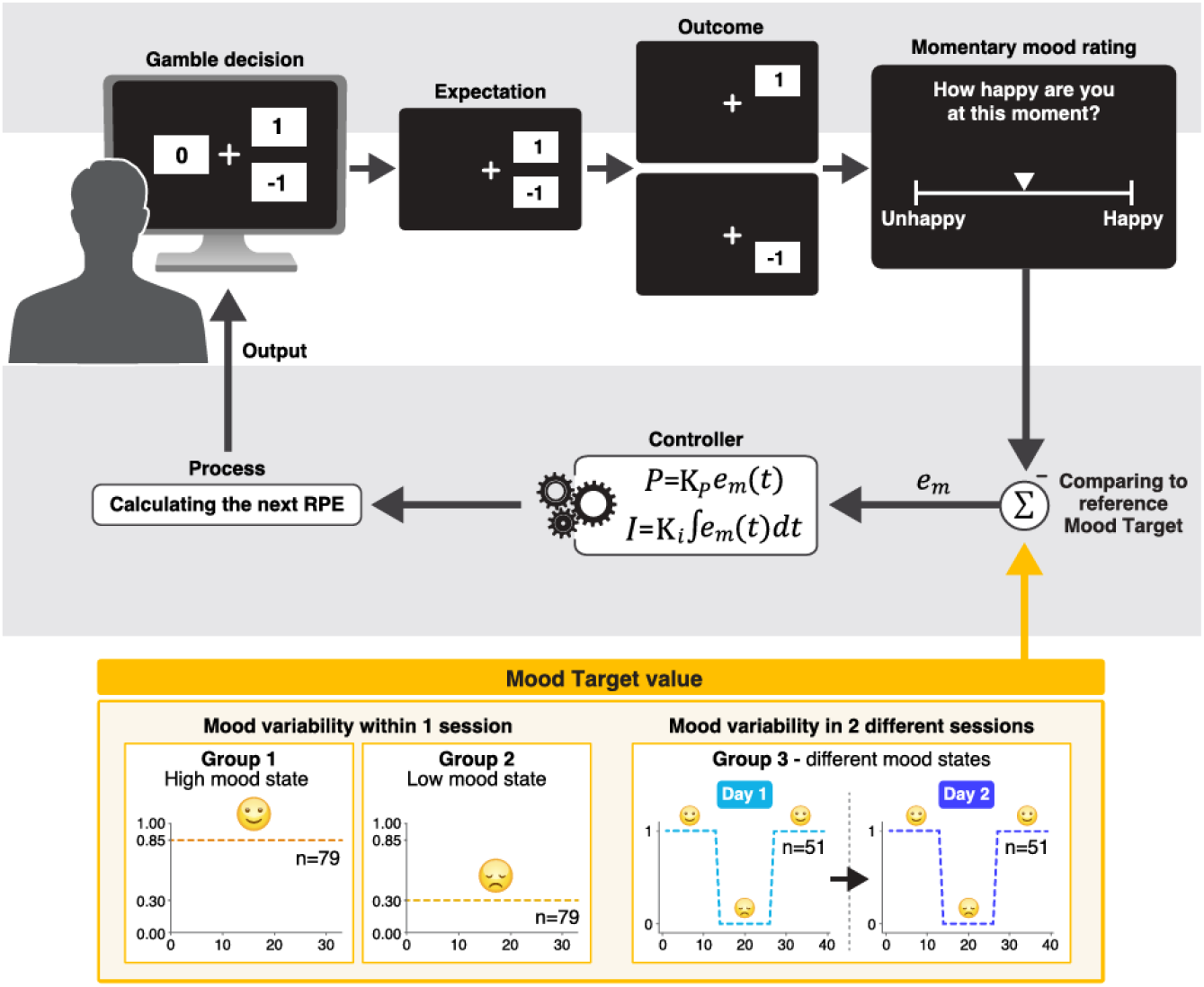
The experimental design. The circuit of the closed-loop mood modulating paradigm is shown, where in each trial, participants first make a gamble decision, choosing between a certain value on the left or a risky gamble on the right. Next, the selected option is shown, followed by the feedback of the win or loss outcome. Every 2-3 trials participants report their momentary mood rating, indicating how happy or unhappy they feel. The difference between this last rating and the mood target (e_m_) is used as input for a proportional-integral (PI) controller, which computes the next Reward Prediction Error (RPE) to move mood toward the target (with a proportional component with gain K_p_ and an integral component with gain K_i_). Bottom section: We used three different types of mood targets in three groups of participants: (1) Single-session experiment: Participants were assigned to either High Mood Target (85%; *n* = 79, Group 1) or Low Mood Target (30%; *n* = 79, Group 2). (2) Two-session test-retest experiment: Each session of the task included three alternating mood targets which the participants repeated on the following day (highest-lowest-highest mood, n=51, Group 3).

We implemented the closed-loop mood modulating paradigm in two different studies: in the first, participants (*n*=158) underwent either a positive or a negative mood modulation toward a mood target of 30% or 85% respectively, and the second was a test-retest study (*n*=51), where participants underwent a three-levels mood modulation (high-low-high) and then repeated the 3-block task on the following day (See Figure 2A and 5A for examples of mood ratings during the different conditions and tasks). Participants provided mood ratings throughout the experiments every 2-3 trials using a slider ranging from “unhappy” to “happy”.

**Figure 2.**
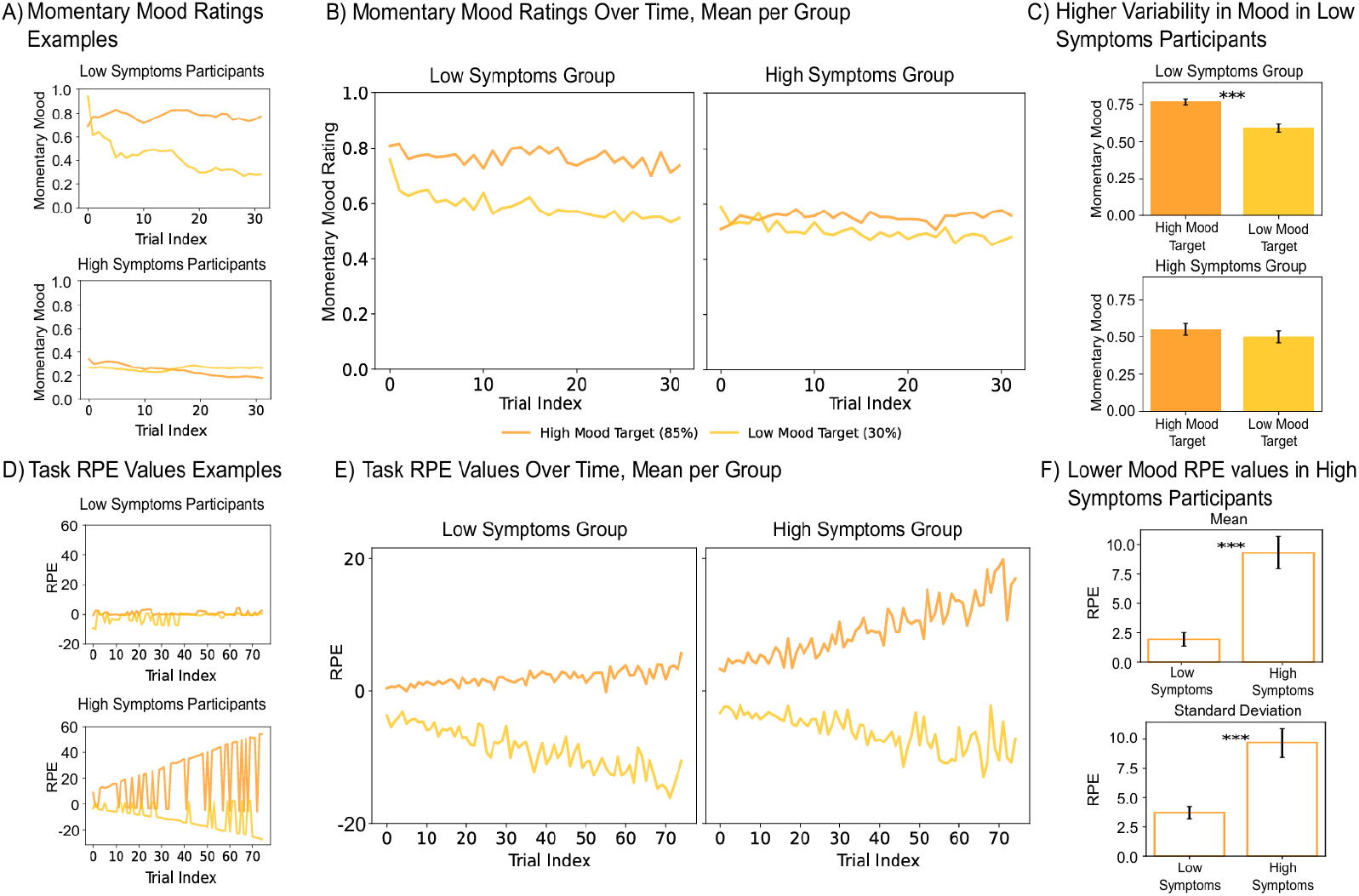
Higher mood variability in low symptoms participants compared to those with higher depressive symptoms. (A) Examples of momentary mood ratings (smoothed with a 5-trial moving average) of one representative participant from each group (the high mood target condition in orange and the low mood target in brighter yellow). (B) Mean momentary mood ratings illustrated per group in each of the conditions. (C) Mood mean by condition in each group. Mood ratings were significantly higher (*p* < 0.001) for the low symptoms group in the High Mood Target condition (n=41) compared to the Low Mood Target condition (n=48) while no significant difference was observed in the high depressive symptoms group. (D) Example of task-generated reward prediction error (RPE) values in each of the two conditions from a representative participant in each group. (E) Mean task RPE values per group. Participants with high depressive symptoms (right) show a marked increase in RPE values over time in the High Mood Target condition, suggesting that the algorithm required greater “effort” to influence mood in this group. (F) Mean and SD of RPE values between groups in the High Mood Target condition (n=79).

In the first low Mood Target condition, we observed a significant negative time effect (*p* < 0.001), where overall participants’ mood declined and stabilized at lower mood levels (suppl. Tabel S1). The extent of mood decline during the task was correlated with depressive symptoms severity, such that the lower the symptoms the stronger the mood decline (more negative mood change) in accordance with the task manipulation (*Pearson r* = 0.25, *p* = 0.024 between CESD-10 vs. last-first three mood ratings difference). To test for a floor-effect, where participants with higher depressive symptoms may not show mood decrease due their lower baseline level, we confirmed that the correlation remains significant after excluding participants with a mean mood lower than 0.2 (*Pearson r* = 0.24, *p* = 0.037). We also found a steeper mood decline in participants with low symptoms when splitting participants into groups with higher (n=31) and lower (n=48) depressive symptoms (with an arbitrary cutoff of CESD-10>=10 following Andresen et al., 1994; Figure 2A-C, suppl. Tabel S2).

The values of RPE stimuli provided during this adaptive task reflect how difficult it is to shift mood, such that the lower the mean RPE, the less responsive the participant in his mood downwards. Analysis of the mean RPE values (Figure 2D-F) revealed a significant positive correlation with depression symptoms (*Pearson r* = 0.294, *p* = 0.008).

Thus, when shifting mood downwards, participants with higher depressive symptoms show lower mood variability compared to participants with lower depressive symptoms.

In the High Mood Target condition, we found a significant negative correlation between depressive symptoms and the standard deviation of mood ratings in the second half of the task after the initial mood transition (*Pearson r* = −0.23, *p* = 0.039), and also with momentary mood variability measured with the Mean Absolute Difference (MAD) index (*Pearson r* = −0.23, *p* = 0.043). When splitting to high and low symptomatic groups, we found again a significant increase in mood over time for the low symptoms participants (*n* = 41, *t* = 2.68, *p* < 0.02), but not for participants with high depressive symptoms (*n* = 38, *t* = −0.99, *p* = 0.33). This suggests that individuals who had a higher initial mood (a Welch independent t-test between initial mood of the two groups with *n* = 79, *t* = −5.6, *p* < 0.001), still shifted their mood upward more strongly (Figure 2C).

Depression scores were also positively correlated with the mean RPE values (*Pearson r* = 0.55, *p* < 0.001), indicating that participants with higher depressive symptoms required greater RPEs to drive mood upward and that while the algorithm increased the RPE values for the depressive individuals, their mood remained unresponsive (Figure 2D-F).

### Computational parameters of mood variability and stability

We employed the Primacy Mood Model, a well-validated model of momentary mood updating in reward-based tasks, an extension of the Happiness Model (Rutledge et al., 2014). This model dissociates the influence of most recent events (Recency coefficient *β*_*R*_) and the influence of early events (Primacy coefficient *β*_*P*_) on mood. This model has shown high accuracy in modeling mood ratings in both adaptive and non-adaptive task designs, in both healthy and clinically depressed participants, and in different age groups (Keren et al., 2021). It also shows high fit to participants’ mood ratings in the current datasets (see examples in suppl. Figure S1), with an average mean squared error (MSE) of 0.0151 (*STD* = 0.022) in the High Mood Target condition and 0.021 (*STD* = 0.027) in the Low Mood Target condition. Importantly, the model was fitted post hoc and did not determine task outcome.

We found that the recency coefficient *β*_*R*_ was positively correlated with the standard deviation of mood ratings in the two different task conditions (Low Mood Target: *Pearson r* = 0.36, *p* = 0.001, High Mood Target: *Pearson r* = 0.62, *p* < 0.001), indicating that participants with greater sensitivity to recent outcomes (higher *β*_*R*_) exhibited more variable mood ratings (Figure 3). This result remained consistent also after recalculating the model’s *β*_*R*_ coefficient using the z-scored values of mood ratings and task reward values, z-scored within-participant (in the Low Mood Target: *Pearson r* = 0.31, *p* = 0.004, and in the High Mood Target: *Pearson r* = 0.43, *p* < 0.0001.) Moreover, the recency coefficient *β*_*R*_ was positively correlated with the range of mood ratings in the two different task conditions, both before and after within-participant z-scoring of the values used in the model (*r* = 0.26, *p* = 0.01 in the Low Mood Target, and *r* = 0.43, *p* < 0.0001 in the High Mood Target, after z-scoring). Thus, participants with greater sensitivity to recent outcomes (higher *β*_*R*_) exhibited a wider range of mood ratings.

**Figure 3.**
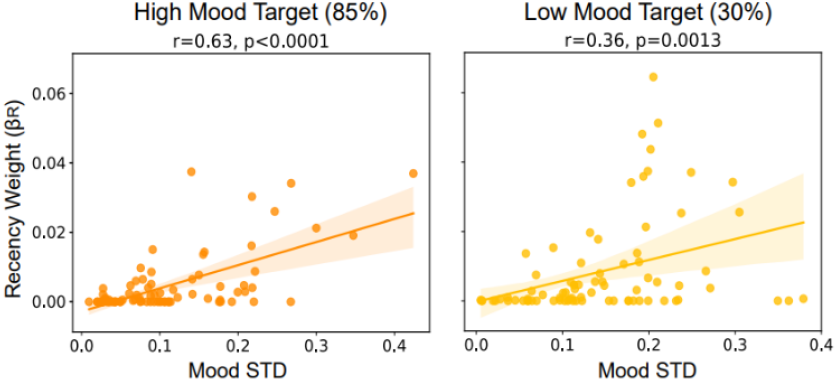
The *β*_*R*_ coefficient is correlated to higher mood variability. Correlations between *β*_*R*_ and standard deviation of mood ratings of each participant, in the two mood target conditions. Each dot represents one participant, and shaded areas represent the 95% confidence intervals of the regression lines.

To examine whether the associations between mood variability and *β*_*R*_ are dominated by baseline mood or the overall mean mood of participants, we conducted a regression analysis including the parameters first mood rating, mean mood, and mood intercept in addition to mood variability. The regression model for both task conditions showed that mood variability had the strongest interaction with *β*_*R*_ relative to the other parameters (*F*(3, 75) = 19.44, *p* < 0.001, explaining 43.7% of the variance in *β*_*R*_ (*R*^2^ = 0.44) in the high mood target, and *F*(3, 75) = 11.37, *p* < 0.001, explaining 31.3% of the variance (*R*^2^ = 0.31) in the low mood target; suppl. Tabel S3). These results were consistent also after z-scoring task values prior to fitting the model in both task conditions.

In the high mood condition, mood variability was also significantly associated with the difference between *β*_*R*_ and *β*_*P*_, such that participants whose *β*_*R*_ had a more dominant influence on mood relative to *β*_*P*_, showed greater variability in mood ratings (*Pearson r* = 0.40, *p* < 0.001). The difference between *β*_*R*_ and *β*_*P*_ was also negatively associated with depression symptoms severity (CESD-10 score), such that individuals with higher *β*_*R*_ relative to *β*_*P*_ had lower depressive symptoms (*Pearson r* = −0.31, *p* < 0.001).

Depression CESD-10 scores were also negatively correlated with the recency influence *β*_*R*_ by itself (*Pearson r* = −0.28, *p* = 0.01; Figure 4), indicating that lower sensitivity to recent outcomes was related to higher depressive symptoms, while importantly, *β*_*P*_ showed the opposite relation – a positive correlation with depression CESD-10 scores (*Pearson r* = 0.28, *p* = 0.01; Figure 4). Thus, individuals with higher depressive symptoms are more influenced by early experiences and show reduced emotional updating and greater rigidity in mood responses, even though the task created a stronger change in RPE values for them (Figure 2F).

**Figure 4.**
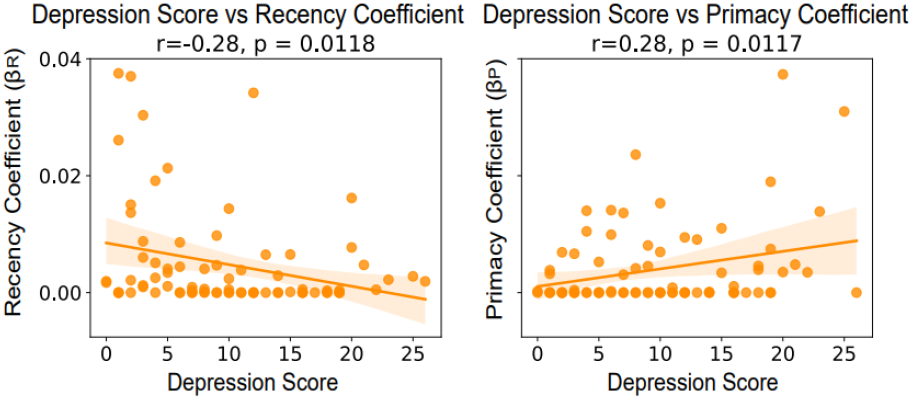
Depression symptoms are negatively correlated with the *β*_*R*_ coefficient and positively correlated with the *β*_*P*_ coefficient. Each plot shows depression scores (CESD-10 questionnaire) against the model-derived parameters *β*_*R*_ (*left*) and *β*_*P*_)*right*) in the High Mood Target condition. Each dot represents one participant, and shaded areas represent the 95% confidence intervals of the regression lines.

Altogether, *β*_*R*_ emerges as a parameter shaping variable mood dynamics, including moment- to-moment variability and overall mood transition in response to the task, in parallel to higher well-being and lower depression symptoms.

### Test-retest reliability of mood updating dynamics and its link to well-being

To test the reliability of these finding over time, other participants completed the closed-loop mood modulating task twice across three different mood targets and across two consecutive days (*n* = 51). The task design integrated two different mood states, alternating the mood target between highest-lowest-highest mood in each of the two sessions (Figure 5A).

**Figure 5.**
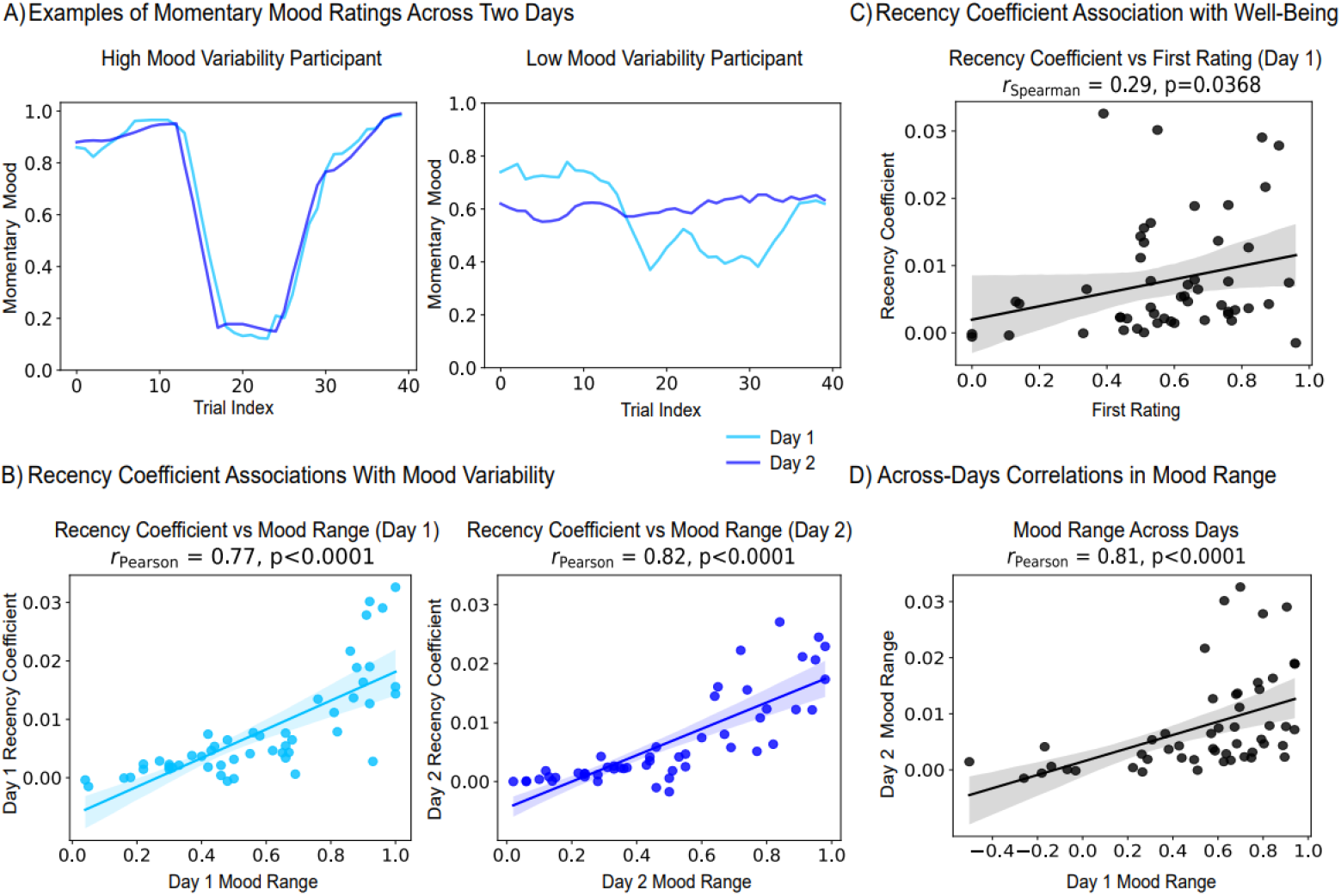
Reliability of emotional updating dynamics and the relation to the recency coefficient and to well-being. (A) Examples of momentary mood ratings of two participants (smoothed with a 5-trial moving average). Left: A participant with high mood range, Right: A participant with lower mood range. (B) Positive Pearson correlations between *β*_*R*_ and mood variability on each of the days (each dot represents one participant and shaded areas represent the 95% confidence intervals around the regression lines). (C) Association between *β*_*R*_ and well-being (see Figure S2 for a demonstration of the connection between well-being and the first momentary mood rating in the task). (D) Across-days correlation between mood range.

Consistent with our single-session findings, also in this separate dataset, *β*_*R*_ was positively correlated with mood range in each of the sessions, reflecting mood variability given that in this task mood was shifted between two extreme mood states (*Pearson r* = 0.77 *in session* 1, *and r* = 0.82 *in session* 2, *p* < 0.001 *for both*). This indicates that participants with higher *β*_*R*_ exhibited greater emotional reactivity and variability during the task (Figure 5B).

Importantly, higher *β*_*R*_ values were also significantly associated with higher well-being, as reflected here by participants’ initial mood rating (*Spearman r* = 0.29, *p* < 0.05; Figure 5C). Finally, we also observed a direct association between well-being and mood variability (the standard deviation of mood ratings correlated with the first mood rating with *Spearman r* = 0.28, *p* < 0.05).

These three correlations form a coherent triangle of interactions between well-being, mood variability, and *β*_*R*_.

Importantly, *β*_*R*_ values of participants were highly correlated across sessions (*Pearson r* = 0.85, *p* < 0.001) and also highly reliable (*ICC* = 0.837, *p* < 0.001), highlighting the within-person consistency of this coefficient and supporting its role as a computational marker. Interestingly, participants also maintained a consistent level of mood variability across days, with high between-session Pearson correlations and test-retest reliability for mood range (*Pearson r* = 0.81, *p* < 0.001, *ICC* = 0.77, *p* < 0.001), indicating an invariant trait-like mood updating dynamics pattern (Figure 5D).

## Discussion

Whether mental health relies more on stability, which buffers mood against fluctuations, or on variability, which enables adaptive responsiveness, is a central question for understanding human well-being and the mechanisms underlying depression. To address this question, we used a twofold computational approach: a closed-loop adaptive task that experimentally induces mood transitions in a personalized manner, and a computational model of mood, allowing to quantify individual mood updating dynamics.

Individuals with higher depression scores showed lower mood variability when mood was modulated both upward and downward, despite receiving higher reward prediction error (RPE) values during the task. This pattern suggests a higher threshold for affective change, where despite the increased stimulus strength, their mood responses remained inert, consistent with emotional rigidity. These findings align with prior evidence linking depression to blunted emotional responsivity, impaired mood regulation, and diminished reactivity to both positive and negative stimuli (Rottenberg et al., 2005; Joormann & Stanton, 2016; Keren et al., 2018, LeMoult & Gotlib, 2019), as well as theories which propose that depression involves excessive reliance on prior expectations and underweighting of novel emotional inputs (Barrett & Simmons, 2015; Barrett et al., 2016; Grahek et al., 2019). These theories are also congruent with the view of depression as a psychopathology of temporal desynchronization, where the past remains dominant and hinders emotional updating (Fuchs, 2001;2013). Interestingly, we find this pattern in participants with higher scores of depression symptoms but without formal diagnosis of depression (can be considered as participants at-risk for clinical depression).

To formulize individual differences in these dynamics, we used the Primacy Mood Model (Keren et al., 2021), which models the influence of recent events versus early outcomes on momentary mood. While Keren et al. (2021) validated the Primacy Mood Model across task conditions and populations and demonstrated its primacy and recency coefficients, the present work focuses on individual differences in its parameters and their links to mood variability and depressive symptoms. We found that higher *β*_*R*_ values, i.e., higher sensitivity of mood to recent outcomes, index greater mood variability and lower depression scores. This links emotional adaptability to resilience and improved mental health (Tugade & Fredrickson, 2004; Kuppens et al., 2010; Waugh et al., 2011; Houben et al., 2015). In contrast, lower *β*_*R*_ values as well as higher *β*_*P*_ values characterized individuals with lower mood variability and elevated depressive symptoms, reflecting diminished incorporation of recent outcomes and anchoring to early events and the maladaptive inertia (Kuppens et al., 2010; Koval et al., 2013; LeMoult & Gotlib, 2019).

A key strength of this study lies in its integration of advanced behavioral and computational approaches, where the closed-loop task allowed to experimentally steer mood in a personalized manner, lowering confounds such as emotional habituation (Jangraw et al., 2023) and enhancing validity by embedding mood ratings in a relational and dynamic decision-making. Embedded partial randomization prevented fixed expectations and reduced demand characteristics. Including complementary experiments – spanning different conditions and timescales – supports the reproducibility and generalizability of our findings.

Several limitations should be noted. First, mood was assessed with a unidimensional valence self-report scale. While clinically relevant (Keyes, 2005; Rottenberg et al., 2005; Rutledge et al., 2014), it does not capture the full complexity of affective states. Future studies could extend this work and explore multidimensional affective states. Second, our two-day test-retest design does not address longer-term resilience or clinical trajectories. Longitudinal work will be helpful in determining whether *β*_*R*_ prospectively forecasts affective outcomes. Finally, although our sample included subclinical variation, replication in clinically diagnosed populations will be informative.

Overall, our findings highlight adaptive emotional variability in response to recent experiences as key to higher well-being and *β*_*R*_ as a candidate computational marker of well-being. Yet, it might be the interplay – rather than tradeoff – between variability and stability that reflects long-term emotional resilience. Notably, our findings provide measurable computational markers detected using a brief behavioral task, thus highlighting the potential of using these markers for early detection of depression and of therapeutic approaches aimed at restoring the temporal balance between past and present experiences.

## Methods

### Participants

#### Single-session participants

Participants were recruited through Prolific (www.prolific.com), an online platform designed for academic and market research, which allows researchers to set customizable eligibility criteria, thereby facilitating high-quality and reliable data collection for scientific studies (Palan & Schitter, 2018). Eligibility criteria for this study included being between 18 and 60 years old, residing in the United States, having no prior exposure to the current experimental task or similar tasks from our lab, and maintaining a Prolific approval rate between 60% and 100%, reflecting a history of successful participation in previous studies.

All participants provided informed consent electronically after reading a detailed description of the research procedures. The study protocol was approved by the Institutional Review Board (IRB) of Bar-Ilan University. Upon completion of the experimental task, participants answered a demographic questionnaire and a set of eight multiple-choice items questions designed to assess their subjective experience and perceived task performance and to report any technical issues. To ensure data integrity, an attentiveness check was placed between two of the items. All participants received compensation through Prolific (£9/hour). In addition, to the standard payment, participants meeting study criteria received a performance-based bonus calculated from their gambling outcomes (£14.38/hour for the High Mood Target and £13.13/hour for the Low Mood Target).

A total of 206 participants completed the study across two experimental conditions. Of these participants, 16 participants with incomplete data files, mostly because they exited the task prematurely, were excluded from analysis. 32 participants were excluded because they failed to provide responses to at least 70% of the mood rating prompts or did not make any choice (gamble or certain) in at least 70% of the trials. Final sample of the single-session experiments thus included 158 participants. For the detailed information of sample size, age and gender of the participants in each of the different task conditions please refer to suppl. Tabel S4.

To assess differences related to depression, all participants completed the Center for Epidemiologic Studies Depression Scale – 10-item version (CESD-10) prior to the task. The CESD-10 is a screening tool used to measure depression risk in the general population, with demonstrated strong validity and reliability (Irwin et al., 1999). Based on the CESD-10 scores, participants were categorized as either with high depressive symptoms (CESD-10 ≥ 10), or with low depressive symptoms (CESD-10 < 10; Andresen et al., 1994).

#### Two-session participants

The data for the two-session analysis in our study were obtained from the open-access dataset accompanying Jangraw et al., 2023, available on the Open Science Framework (https://osf.io/km69z/). From this dataset, we included participants who completed the closed-loop version of the task in two consecutive sessions.

Participants were recruited via Amazon Mechanical Turk (Amazon.com, Inc., Seattle, WA), (Paolacci et al., 2010). Eligibility criteria for this study required participants to be at least 18 years old, reside in the United States, have no prior exposure to the current experimental task, have completed over 5,000 tasks for other requesters, and have over 97% of their tasks accepted as satisfactory to the requester. These criteria promise better validity of the data and are congruent with recommendations for online data collection (Paolacci et al., 2010; Peer et al., 2014; Chandler & Shapiro, 2016).

All participants received identical written instructions and provided informed consent on a web page where they were required to click ‘I Agree’ to participate. Since this study did not engage in direct intervention or interaction with participants, nor did they collect any personally identifiable private information, the study was exempt from institutional review board (IRB) oversight, as determined by the National Institutes of Health (NIH) Office of Human Subjects Research Protections. The consent procedure and details of the task and survey were reviewed and approved by the Office of Human Subjects Research Protections.

All participants received compensation. In addition to the standard payment, participants meeting study criteria received a performance-based bonus calculated from their gambling outcomes.

To be included in the dataset, participants were required to complete both the task and the associated survey within a 90-minute period, beginning at the time they accepted the assignment. Completion was verified by submitting a six- to ten-digit code presented at the end of each component (task and survey). While the second session could be completed at any time within the following 3 days, both the task and the survey for that session had to be completed within the same 90-minute window. The mean test-retest interval was 1.05 days (STD = 0.20).

Of the 66 individuals who completed the task on the first session, 53 (80.3%) completed the second session. Two participants had incomplete data files and were excluded. The final sample was 51 participants. For the detailed information on sample size, age and gender of the participants please refer to suppl. Tabel S4.

### Experimental Design

All tasks were developed using PsychoPy (version 2023.2.3 for the single-session task and version 2020.1.2 for the two-session task) and deployed via Pavlovia, an online behavioral experiments platform. Pavlovia uses the JavaScript-based PsychoJS library to present experiments and integrates with the Prolific and Amazon Mechanical Turk recruitment platforms.

### Closed-loop mood modulation task

The experimental task was adapted from a probabilistic reward paradigm used in previous studies (e.g., Rutledge et al., 2014; Keren et al., 2021). It was composed of choice trials where participants had to make a quick decision whether to gamble or take a certain value of points, interleaved with repeated ratings of momentary mood (Figure 1).

Each trial in the gambling task began with a decision screen with three boxes (the decision phase, shown for 3 seconds). The left box showed a certain value, while the two boxes on the right-side presented two potential outcomes, one higher than the other. These values represented game points and participants were instructed that winning more points will lead to receiving a higher monetary bonus for the task (ranging between $0-6).

Participants chose whether to gamble or accept the certain value by pressing the right or left arrow key, with default gambling if no choice was made. Upon selection, the unchosen option disappeared, and if a gamble was made, both optional values were shown (expectation phase, 2-4 seconds). The outcome was then displayed (outcome phase, 1 second). Participants rated their momentary mood every 2-3 such trials by answering, *“How happy are you at the moment?”*. First, the mood question appeared on the screen without the option to rate mood (mood question phase, 1.5-3 seconds), followed by the appearance of the mood slider and the option to rate mood (mood rating phase, 4-4.5 seconds). Participants adjusted their mood rating using the left/right arrow keys, pressed continuously for a quicker change, and confirmed their choice with the spacebar.

***The values for gambling outcomes*** were determined based on the last mood rating of individuals such that a certain RPE value was reached.

The next RPE was calculated using a Proportional-Integral (PI) control algorithm, based on the discrepancy between the participant’s reported mood and the target mood. The RPE reflected a weighted combination of the proportional mood error (the difference between last mood rating and target mood) and the cumulative error (sum of the proportional errors from the beginning of the block). A minimum RPE of ±0.03 was enforced to ensure meaningful feedback.

The RPE was calculated according to the following equation:

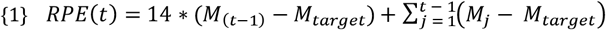

Where *t* represents the trial number, *M*_(*t*)_ denotes the mood reported following trial *t*, and *M*_*target*_ indicates the target mood in that block.

Overall RPE was calculated to adjust mood toward the target: providing positive RPEs when mood was below the target and negative RPEs when mood was above it.

To maintain unpredictability and engagement, 30-35% of the trials were set to be *incongruent* trials, where RPEs opposed the target direction and were downscaled to reduce their mood influence, according to the following equation:

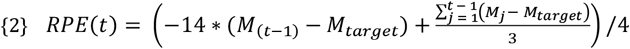

The next trial gambling values were determined according to the RPE value that was calculated. First, a base value was randomly selected between -4 and 4, with possible values increasing in steps of 0.1, and the alternative gambling value was calculated by adding or subtracting the calculated RPE value. The “certain” value was computed as either the mean between the two values or as the (higher value + 2 x lower value) / 3, designed to incentivize gambling.

***In the single-session experiments***, each participant completed 75 trials, the experimental procedure lasted approximately 25 minutes and consisted of several distinct phases. Following informed consent, participants completed the CESD-10 questionnaire. Participants then read the first part of the task instructions and provided a baseline general mood rating. After reading the second part of the instructions, participants completed a brief gambling task training session to ensure comprehension. Subsequently, they engaged in the main task, which included interleaved momentary mood ratings. Finally, participants completed a survey. The task aimed to stabilize participants’ mood at one of two predefined target levels: (1) In the *High Mood Target* condition, the goal was to elevate and maintain mood at a high positive level, targeting 85% on the mood scale; (2) In the *Low Mood Target* condition, the goal was to lower and stabilize mood at a reduced level, targeting 30% on the scale.

Participants provided a total of 33 mood ratings, including the baseline rating before the first trial. If a participant failed to provide a mood rating, the most recent available rating was used for task values calculations. In addition, for data analysis, missing mood values were linearly interpolated to create a continuous mood time series per participant.

Trial congruency varied by condition, with 35% of trials set as incongruent. Moreover, in these versions, the absolute value of the mood error was smaller than 0.04, the value of the sum of the errors was reset to zero to prevent RPE escalation when participants are close to the mood target. When the mood rating equaled the target value, values of the presented options were randomized ±0.1 around the base value.

***In the test-retest two-session experiments***, each of the sessions lasted approximately 17.3 minutes and consisted of several distinct phases, including task instructions, providing baseline mood rating and subsequently, the main gambling task.

The mood target for the controller during each of the sessions was set to: maximal mood value of the scale of 100% (32 trials, 13 mood ratings), minimal mood value of 0 (32 trials, 13 mood ratings), maximal mood value again (32 trials, 14 mood ratings).

30% of the trials were set as incongruent trials. In each session, participants provided a total of 40 mood ratings.

### Data Analysis

**Within-session mood flexibility** describes how much individual’s mood changes during a single experimental session, reflecting their emotional responsiveness to the task. We assessed this first using a linear mixed-effects (LME) model. Analyses were performed using JASP (version 0.19.3.0), an open-source statistical software. Mood rating times were converted to minutes to meet the model’s convergence requirements while preserving interpretability. The fixed effects included Time (*t*), Group, and their interaction (Time × Group), with mood ratings (*M*) as the dependent variable. To account for individual differences in baseline mood and temporal trends, random intercepts and slopes for Time were included for each participant. The model was specified as:

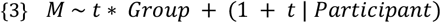

In addition to the Linear Mixed-Effects models, we conducted repeated-measures ANOVA to further examine whether mood ratings changed significantly during the task. Specifically, for each condition (High Mood Target and Low Mood Target) we used the time points of mood ratings as a within-subject factor to assess temporal trends in mood.

Paired-sample t-tests were performed to assess mood changes within a session by comparing the mean of the last three mood ratings with the mean of the first three ratings.

In the test-retest two session experiments, mood flexibility was assessed with the range of mood, i.e., the difference between highest and lowest mood ratings.

**Across-session mood dynamics consistency and reliability** refer to the extent to which an individual exhibits similar mood updating patterns across the two experimental sessions conducted on different days. To evaluate this, we employed two different measures: (1) A Pearson correlation of the mood range between sessions. This measure allowed us to assess whether individuals maintained a similar mood variability level across days. (2) The Inter-Class-Correlation (ICC2), calculated for both *β*_*R*_ coefficient and the mood range, to quantify the test-retest reliability of these indicators of emotional updating and variability across sessions.

Together, these indices capture consistency in emotional dynamics such as emotional range, and the temporal structure.

### Computational model – the Primacy Mood Model

The Primacy Mood Model integrates two key components formulated as the following equation:

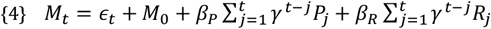

*M*_0_ is the mood intercept, *ϵ*_*t*_ is a random noise, and *γ* is exponential discounting factor. *P*_*t*_ is the Primacy influence term:

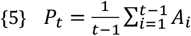

Where all previous outcomes (*A*_*i*_) are averaged to describe a history-based influence on mood, which results in a primacy weighted cumulative influence of outcomes. And respectively, the *β*_*P*_ is the individual coefficient of the primacy influence on mood. *R*_*t*_ is the Recency influence term:

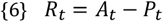

By subtracting the last outcome from the average of all previous outcomes, we reach an RPE-like term, which represents the influence of the difference between these different temporal weightings of outcomes. Respectively, *β*_*R*_ is the individual coefficient of the recency influence on mood.

Initial parameter values were set as follows: *M*_0_ = 0.5, *γ* = 0.8, *β*_*P*_= 0.01, *β*_*R*_ = 0.005. Parameter bounds were defined to ensure realistic and stable predictions: *M*_0_ ∈ [0.0, 1.0], *γ* ∈ [0.0, 1.0].

Model fitting was performed using L1-regularized absolute error minimization. To prevent overfitting, a regularization penalty of 0.1 was applied to each beta parameter. This penalty was adjustable at runtime, depending on study-specific requirements. Model implementation was conducted in Python and optimization was performed using the “minimize” function from the SciPy optimize module, with constraints applied according to the defined parameter bounds. To evaluate model performance and assess the quality of fit, we computed the Mean Squared Error (MSE) between predicted and actual mood ratings.

### Regression Analysis

We conducted a series of linear regression analyses to explore whether individual differences in the model-derived parameter *β*_*R*_ could be accounted for by participants’ mood characteristics versus the contribution of moment-to-moment mood fluctuations. Specifically, we examined the contribution of three mood-related predictors: (1) the overall mean of mood ratings, (2) mood variability, quantified as the standard deviation (SD), and (3) the initial mood rating reported at the beginning of the task.

Separate models were estimated for each experimental condition with *β*_*R*_ as the dependent variable. This allowed us to assess which mood features best predict participants’ responsiveness to recent outcomes across different regulatory goals.

The regression model was specified as follows {7}:

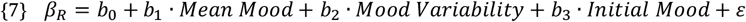

Where *b*_1_, *b*_2_, *b*_3_ are the coefficient weights and *ε* is the residual error.

### Additional Statistical Analyses

We z-scored all the mood ratings values and all gamble outcomes within subject and re-modeled the data to then re-extract the *β*_*R*_ and *β*_*p*_ coefficients. This normalization allowed us to test correlations with depressive symptoms using relative coefficients.

We conducted independent-sample t-tests to compare low symptoms and high symptoms participants across key outcome variables, including average mood, initial mood rating, mood variability, and average RPE values, separately for each condition. In cases where Levene’s test indicated unequal variances between groups, we applied Welch’s t-test, which provides a more reliable estimate under variance heterogeneity.

We further employed Pearson and Spearman correlation analyses to investigate relationships between behavioral measures, model parameters, and clinical indices.

The Mean Absolute Difference (MAD) index was calculated for the second half of the mood ratings, from the difference between each two consecutive mood ratings.

## Data & Code Availability

All data and analysis code supporting the findings of this study are available at the following GitHub repository: https://github.com/NoaN16/EmotionalUpdating.

## Acknowledgements

This study was supported by the Israeli Science Foundation (ISF; grants 3066/24 and 3627/24, to H.K.). We thank Prof. Shimon Marom for introducing us to the theoretical framework that informed parts of the discussion, and Boaz Harel for his technical assistance and support.

## Notes

https://github.com/NoaN16/EmotionalUpdating

### Competing Interest Statement

The authors have declared no competing interest.

