## Supplementary for "Dynamic emotional updating as a computational marker of well-being"

| Condition | Term | Estimate | Std. Error | df | t | p-value |
|---|---|---|---|---|---|---|
| *High Mood Target* (85%) | Intercept | 0.667 | 0.027 | 75.96 | 24.440 | <0.001 |
| | Time | -0.00072 | 0.002 | 75.038 | -0.340 | 0.735 |
| | Group (Low symptoms) | 0.139 | 0.027 | 75.96 | 4.976 | <0.001 |
| | Time ✻ Group | -0.003 | 0.002 | 75.038 | -1.302 | 0.199 |
| *Low Mood Target* (30%) | Intercept | 0.647 | 0.030 | 76.834 | 21.422 | <0.001 |
| | Time | -0.01 | 0.002 | 75.409 | -4.201 | <0.001 |
| | Group (Low symptoms) | 0.061 | 0.030 | 76.834 | 2.007 | 0.048 |
| | Time ✻ Group | -0.003 | 0.002 | 75.409 | -1.309 | 0.194 |
| *Low Mood Target* (30%) | Intercept | 0.599 | 0.053 | 70.217 | 11.218 | <0.001 |
| | Time | -0.006 | 0.003 | 55.597 | -1.722 | 0.091 |
| | Group (Low symptoms) | 0.018 | 0.0053 | 70.217 | 0.338 | 0.736 |
| | Time ✻ Group | 0.001 | 0.003 | 55.597 | 0.312 | 0.756 |

**Table S1.** LME Results Tables.
Fixed-effects estimates from linear mixed-effects models fitted to mood ratings across conditions: (1) *High Mood Target* (85%; $n = 79$; $n_{(low\ symptoms)} = 41$, $n_{(high\ symptoms)} = 38$), and (2) *Low Mood Target* (30%; $n = 79$; $n_{(low\ symptoms)} = 48$, $n_{(high\ symptoms)} = 31$), including analyses for all trials and for trials from 17 onwards. For each model, the table reports the estimated coefficients (β), standard errors (SE), degrees of freedom (*df*), *t*-values, and corresponding *p*-values.

| Time | Group | Estimate | SE | 95% CI | |
|---|---|---|---|---|---|
| | | | | Lower | Upper |
| 5.134 | Low symptoms | 0.642 | 0.031 | 0.581 | 0.703 |
| 11.220 | Low symptoms | 0.565 | 0.031 | 0.503 | 0.626 |
| 17.306 | Low symptoms | 0.488 | 0.040 | 0.409 | 0.567 |
| 5.134 | High symptoms | 0.552 | 0.039 | 0.476 | 0.628 |
| 11.220 | High symptoms | 0.511 | 0.039 | 0.435 | 0.588 |
| 17.306 | High symptoms | 0.471 | 0.050 | 0.373 | 0.569 |

**Table S2.** Estimated Marginal Means of Mood Ratings Across Timepoints for Each Group in the Low Mood Target Condition *(n* = 79; $n_{(low\ symptoms)} = 48$, $n_{(high\ symptoms)} = 31$).

| Predictor | *High Mood Target (85%)* | *Low Mood Target (30%)* |
|---|---|---|

| | | |
|---|---|---|
| Intercept | -0.0087 (p=0.003) | 0.0280 (p<0.0001) |
| Mood Mean | 0.0085 (p=0.131) | -0.0167 (p=0.032) |
| Mood STD | 0.0691 (p<0.0001) | 0.0461 (p=0.008) |
| First Mood Rating | ≈ 0.0000 (p=1) | -0.0142 (p=0.016) |

**Tabel S3.** Linear regression results predicting $\beta_R$ from mood features in each condition.
*Each column presents standardized regression coefficients (β) and associated p-values for predictors.

| | | N | Age (mean ±std) | Gender (F/M) |
|---|---|---|---|---|
| **Single-session** | *Low Mood Target (30%)* | 79 | 36.87±11.49 | 39/40 |
| | *High Mood Target (85%)* | 79 | 39.48±10.73 | 42/37 |
| **Two-sessions** | | 51 | 37.30±10.91 | 23/28 |

**Table S4.** Participant Demographic Information by Experimental Condition.

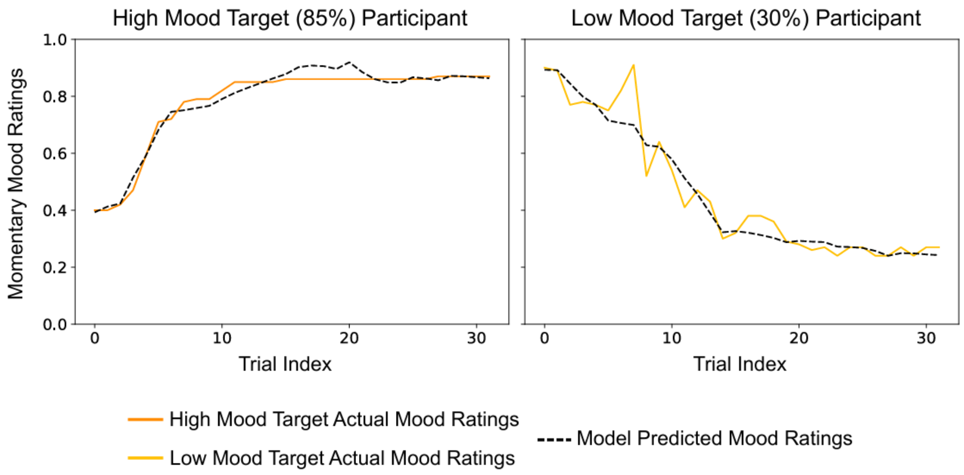


**Figure S1.** Examples of individual participants' mood ratings during the task and the simulated mood according to the model predictions. The left panel presents an example from a participant in the High Mood Target condition, and the right panel shows an example from a participant in the Low Mood Target condition. Solid lines represent participants' actual momentary mood ratings, while dashed black lines represent the mood predictions generated by the Primacy Mood Model.

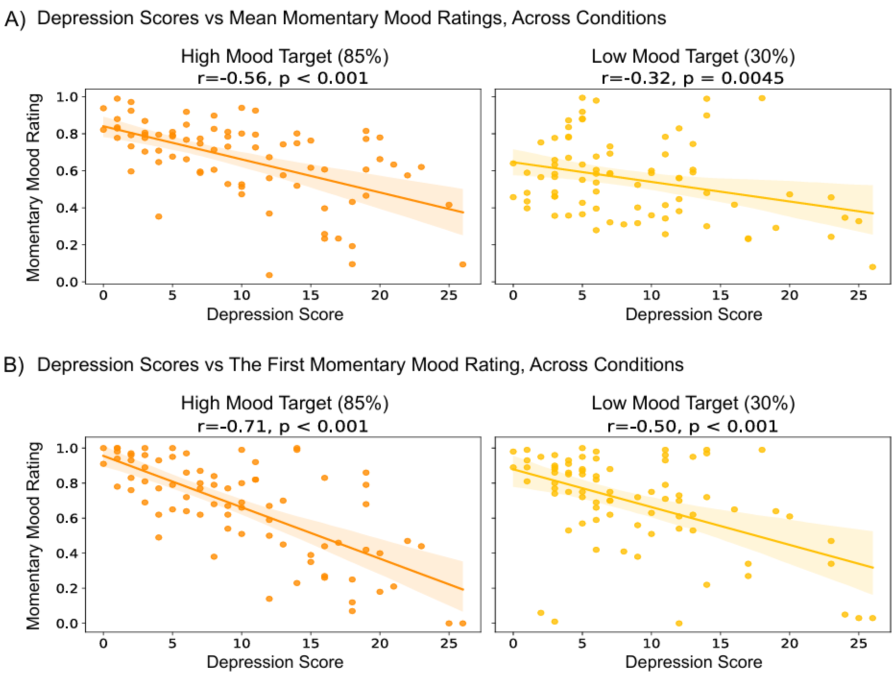


**Figure S2.** Depression score interaction with task momentary mood ratings. All scatterplots show Pearson correlations. Each dot represents one participant, and shaded areas denote the 95% confidence intervals of the regression lines. Left: The High Mood Target condition (*n*=79), right: the Low Mood Target condition (*n*=79). (A) Correlations between Depression scores (CESD-10) and the mean momentary mood ratings. (B) Depression scores were also negatively correlated with the first momentary mood rating in both conditions, showing an even stronger relationship and suggesting that the first mood rating may serve as a more robust indicator of participants' baseline affective state.
